# Sheltered load in fungal mating-type chromosomes revealed by fitness experiments

**DOI:** 10.1101/2024.09.10.612177

**Authors:** Lou Guyot, Elizabeth Chahine, Elsa De Filippo, Christophe Lalanne, Sylvain Brun, Fanny E. Hartmann, Tatiana Giraud

## Abstract

Sex chromosomes and mating-type chromosomes can carry large regions with suppressed recombination. As a result of a lower efficacy of selection, recessive deleterious mutations are expected to accumulate in these non-recombining regions. Multiple genomic analyses have indirectly inferred the presence of deleterious mutations in sex and mating-type chromosomes, but direct experimental evidence remains scarce. Here, we performed fitness assays in fungi with megabase-large and young non-recombining regions around the mating-type locus, using three Sordariales species, to test whether heterokaryons (diploid-like, heterozygous at the mating-type locus) exhibited a fitness advantage over homokaryons (haploid-like, with a single mating-type allele), in terms of spore germination dynamics or mycelium growth speed, under different conditions of light and temperature. We found a faster growth of heterokaryons compared to one of the homokaryons for *Podospora anserina* at 18°C and for *Schizothecium tetrasporum* and *Schizothecium tritetrasporum* at 22°C under light. These findings suggest the presence of a sheltered load, i.e., recessive deleterious mutations at the heterozygous state in or near non-recombining regions, associated to a specific mating-type allele. Genomic analyses indeed suggested that the non-recombining regions around the mating-type locus likely carries heterozygous deleterious mutations, while the rest of the genome was mostly homozygous. We also showed that the difference in growth rates did not result from different numbers or densities of nuclei between homokaryons and heterokaryons. Leveraging the experimental assets of fungi, allowing cultivating separately haploid-like and diploid-like life stages, our experiments provided one of the rare direct experimental evidence of sheltered load around mating-compatibility loci, which is crucial for our understanding of sex-related chromosome evolution.

**Social Media Abstract:** Experimental evidence for sheltered load around the mating-type locus in filamentous fungi: slower growth of haploid versus diploid-like mycelia, revealing recessive deleterious mutations, in *Podospora anserina* and other Sordariales fungi.

## Introduction

Even in sexual species, genomes can display large non-recombining regions (Wright et al., 2016, Schwander et al., 2014, Bergero & Charlesworth, 2009). Such non-recombining regions are particularly frequent around sex-determining loci and mating-type loci (Charlesworth, 2017, Ma & Veltsos, 2021, Hartmann et al., 2021b, Jay et al., 2024, Nicolas et al., 2004, Bachtrog, 2013). Deprived of recombination, these genomic regions experience a reduced efficacy of selection (Felsenstein, 1974, Muller, 1932, Hill & Robertson, 1966, Rice & Chippindale, 2001). As a consequence, it has been theoretically suggested that non-recombining regions are bound to degenerate (Smith & Haigh, 2009, Muller, 1932, Hill & Robertson, 1966, Fisher, 1930), i.e., to accumulate recessive deleterious mutations following recombination suppression. In addition, theoretical models have shown that recessive deleterious mutations can be maintained for some times in populations when tightly linked to self-incompatibility or mating-type loci, especially when weakly deleterious, generating mutational load associated to a specific self-incompatibility or mating-type allele (Tezenas et al., 2023, Antonovics & Abrams, 2004, Glémin et al., 2001, Uyenoyama, 2005). One therefore expects recessive deleterious mutations to be present in or near non-recombining regions around sex-determining or mating-type determining alleles. Such recessive deleterious mutations associated with a permanently heterozygous allele constitute a sheltered load, as they are also heterozygous and therefore not exposed to selection, being protected by the functional copy of the alternative allele in the heterozygote.

The presence of deleterious mutations in non-recombining mating-type and sex chromosomes has been indirectly inferred through genomic analyses, that have for example revealed gene disruptions, gene losses, non-synonymous substitutions, transposable element accumulation and chromosomal rearrangements (Carpentier et al., 2022, Vittorelli et al., 2023, Duhamel et al., 2023, Bachtrog, 2013, Hough et al., 2014, Whittle et al., 2011). However, there has been little direct evidence for the existence of such sheltered load through experimental fitness assays so far. To perform such experiments, one indeed needs to compare fitness between a haploid phase or a homozygous state and the heterozygous diploid state, i.e., between situations where mutations are exposed to selection to situations where deleterious mutations are sheltered at a heterozygous state. While Y-like chromosomes are never homozygous, non-recombining regions can be found homozygous in some autosomal supergenes, i.e. large regions in autosomes governing adaptive and complex phenotypes and segregating as a single unit (Schwander et al., 2014). Using such autosomal supergenes that can be homozygous, the existence of sheltered load could be demonstrated in some cases (Jay et al., 2021). A few balanced haplo-lethal systems even exist, in which recessive deleterious mutations are lethal at the homozygous state in non-recombining supergenes (Wielstra, 2020, Hill et al., 2023). However, in the majority of organisms with non-recombining regions on sex-related chromosomes, haploid or homozygous states do not exist in the life cycle and cannot be generated. The rare exceptions include self-incompatibility loci in plants that can be rendered homozygous under certain conditions. Sheltered load was found associated with some self-incompatibility alleles in the plants *Arabidopsis halleri* (Stone, 2004, Llaurens et al., 2009, Le Veve et al., 2024) and *Solanum carolinense* (Mena-Alí et al., 2009).

Fungi can be highly tractable models for investigating the existence of a sheltered load in sex-related chromosomes. Indeed, there can be recombination suppression in large genomic regions around their mating-type loci in species with a long heterokaryotic (diploid-like) phase, i.e., with nuclei of opposite mating types per cell, but unfused, which is functionally similar to a diploid stage (Branco et al., 2018, Hartmann et al., 2021b, Grognet et al., 2014, Menkis et al., 2008, Vittorelli et al., 2023). Genomic footprints of degeneration have been found in these non-recombining regions, in the form of gene losses, transposable element accumulation, non-synonymous substitutions and non-optimal codon usage and expression (Carpentier et al., 2022, Vittorelli et al., 2023, Duhamel et al., 2023, Whittle et al., 2011).

Mating types in fungi do not prevent selfing, as mating compatibility is controlled at the haploid stage (Billiard et al., 2012, Giraud et al., 2008). As a matter of fact, the fungal lineages with recombination suppression around their mating-type loci most often undergo regular automixis i.e., intra-tetrad selfing (Billiard et al., 2012, Giraud et al., 2008, Hood & Antonovics, 2000, Zakharov, 2005, Zakharov, 2023, Grognet et al., 2014, Jay et al., 2024, Vittorelli et al., 2023). Such life cycles in fungi are called pseudo-homothallic, with the production of self-fertile heterokaryotic spores, encompassing two nuclei of opposite mating types originating from a single tetrad after meiosis, thus germinating as a self-fertile heterokaryon and undergoing mostly automixis. Self-sterile spores with a single mating type, germinating as homokaryotic mycelia, can also generally be produced in laboratory conditions, although their importance in nature is unknown.

With such life cycles, experiments can be set up for comparing fitness between homokaryons (i.e., with a single type of nuclei per cell) and heterokaryons (i.e., with nuclei of opposite mating types in each cell) in order to study the heterozygous advantage in the region around the mating-type locus. Indeed, natural heterokaryotic strains are typically highly homozygous genome-wide except around the mating-type locus, because of high selfing rates (Branco et al., 2017, Grognet et al., 2014, Hartmann et al., 2021a, Hartmann et al., 2021b, Vittorelli et al., 2023). Therefore, any variability in fitness between homokaryotic and heterokaryotic stages should mainly be due to heterozygous recessive deleterious mutations present in the non-recombining region around the mating-type locus and possibly in its flanking region due to linkage disequilibrium (Branco et al., 2017, Tezenas et al., 2023, Glémin et al., 2001) (Fig 1A). Recessive lethal alleles linked to the mating-type locus have been documented in anther-smut fungi of the *Microbotryum* genus and in the fungal-like oomycete *Phytophthora infestans* (Judelson et al., 1995, Hood & Antonovics, 2000, Thomas et al., 2003, Oudemans et al., 1998).

**Figure 1:**
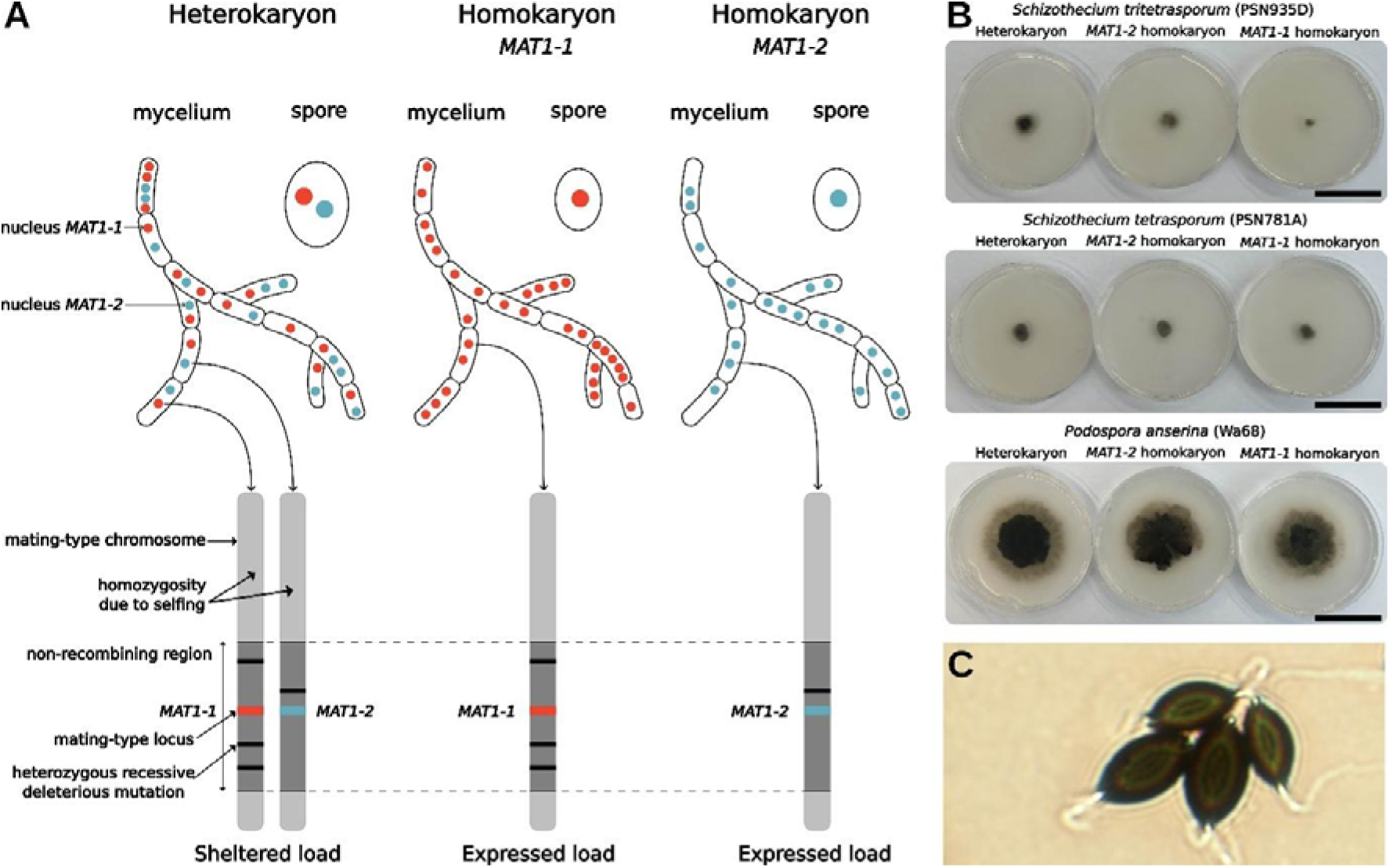
Fitness experiments to test for the presence of a sheltered load in pseudo-homothallic ascomycete fungi. **A)** Schema illustrating how the life cycle of pseudo-homothallic fungi can be used to test for the existence of sheltered load (i.e. the presence of heterozygous recessive deleterious mutations) in the non-recombining region around the mating-type locus. Pseudo-homothallic ascomycete fungi produce heterokaryotic spores (with nuclei of opposite mating types), and with therefore a main phase in their life cycle that is heterokaryotic, i.e. diploid-like. **B)** Pictures of mycelium after six days of growth on Petri dishes under light conditions at 22°C, with heterokaryons at left, *MAT1-2* homokaryons in the middle and *MAT1-1* homokaryons at right, for *Schizothecium tritetrasporum* (strain PSN935D), *Schizothecium tetrasporum* (strain PSN781A) and *Podospora anserina* (strain Wa68), from top to bottom. Scale bar: 4 cm. **C)** Picture of *Schizothecium tetrasporum* spores (strain CBS815.71), with two small spores on the top right and two large spores on the bottom left. The large spores have length and width of ca. 25 x 12 µm.

We studied here three species of two phylogenetically distant fungal species complex within the Sordariales order of Ascomycota: *Schizothecium tetrasporum* and *Schizothecium tritetrasporum,* belonging to the *Schizothecium tetrasporum* species complex (Vittorelli et al., 2023, De Filippo et al., 2025), and *Podospora anserina* (Grognet et al., 2014, Hartmann et al., 2021a, Boucher et al., 2017). These species are coprophilous, saprophyte or endophyte, and can be found in herbivore faeces and soil (Silar, 2020). These fungi are excellent biological models to investigate the presence of a sheltered load around their mating-type locus, displaying two mating-type alleles (*MAT1-1* and *MAT1-2*, also called *mat-* and *mat*+, respectively). Indeed, these fungi display recombination suppression around their mating-type locus, as shown by progeny segregation and nucleotide divergence analyses (Grognet et al., 2014, Hartmann et al., 2021a, Vittorelli et al., 2023). These species are pseudo-homothallic, i.e. with their main life cycle phase being heterokaryotic (Grognet et al., 2014, Vittorelli et al., 2023, De Filippo et al., 2025), so that sheltered load can be expected. Recombination suppression around the mating-type locus spans approximately 1 Mb in the *S. tetrasporum* and *P. anserina* species complexes and has been suggested to be quite recent, as indicated by the low divergence between mating types and the lack of trans-specific polymorphism (Vittorelli et al., 2023, Grognet et al., 2014, Hartmann et al., 2021a). Indirect evidence of sheltered load has been reported by genomic analyses in the non-recombining regions of the reference strains, in particular a few gene losses or disruptions (Vittorelli et al., 2023, Grognet et al., 2014). Furthermore, recombination suppression around the mating-type locus represents convergent events between the *S. tetrasporum* and *P. anserina* species complexes, that diverged over 150 million years ago (Hyde et al., 2017), which provides independent events of recombination suppression to study. Even within each of these two species complexes, recombination suppression has likely evolved independently in the various species, as suggested by the lack of trans-specific polymorphism and sizes of the non-recombining regions (Hartmann et al., 2021a). In addition, natural strains are highly homozygous genome-wide, except around the mating-type locus, as shown in multiple strains of *P. anserina* and in the reference strain of *S. tetrasporum* (Vittorelli et al., 2023, Grognet et al., 2014, Hartmann et al., 2021a). Another asset of these models is that one can cultivate *in vitro* homokaryotic and heterokaryotic mycelia (Fig 1B). In addition, while most ascospores are heterokaryotic, homokaryotic spores are sometimes produced and are easily distinguishable based on their smaller size (Vittorelli et al., 2023, Silar, 2020) (Fig 1C). In *S. tetrasporum* and *P. anserina,* previous experiments on a single strain each did not detect any heterokaryon advantage (Grognet et al., 2014, Vittorelli et al., 2023). Studying multiple strains is however essential in evolutionary biology, as the effects of deleterious mutations can depend on epistatic effects and/or be present in only some individuals in populations, while still having important population effects. Studying various situations and diverse fitness components is also important, as the sheltered load may only be detectable in some conditions, for example if recessive deleterious mutations affect genes expressed only in specific situations, or if stressful conditions exacerbate deleterious effects. Examples of stressful conditions for these species can be temperature or darkness, as fungi do not tolerate high temperatures and use light as an important regulator of diverse physiological processes, as shown in particular in *P. anserina* (Shen et al., 2022, Silar, 2020).

Here, we thus conducted assays using different fitness proxies on a dozen strains of each of *S. tetrasporum*, *S. tritetrasporum* and *P. anserina*, by comparing heterokaryons and homokaryons, in terms of mycelium growth speed under light or dark, at different temperatures, and in terms of spore germination dynamics, and we analysed genomes. We expected that i) strains would be homozygous genome-wide except around the mating-type locus, ii) heterozygous non-synonymous mutations would be present in the non-recombining region around the mating-type locus of our strains, iii) at least one homokaryon would have a lower fitness than the heterokaryon, at least under some conditions, due to recessive deleterious alleles associated to a specific mating type. Our experiments revealed as expected a fitness advantage of heterokaryons over one of the homokaryons (either *MAT1-1* or *MAT1-2*), in at least one tested condition, for *S. tetrasporum*, *S. tritetrasporum* and *P. anserina*. This indicates the likely existence of a sheltered load, associated to a specific mating type allele, in the non-recombining region present around the mating-type locus and/or in its flanking region. Genome analyses indeed showed that most of the strains used here in *S. tetrasporum* and *S. tritetrasporum* were homozygous genome-wide, except around the mating-type locus, as previously shown in multiple strains in *P. anserina* (Hartmann et al., 2021a) and in the reference strain of *S. tetrasporum* (Vittorelli et al., 2023). Furthermore, we detected heterozygous non-synonymous mutations in the non-recombining region, with in particular gains of stop codons and loss of start codons, that are likely deleterious. Counts of nuclei in the reference strain of *P. anserina* suggested that the difference in growth rates did not result from different numbers or densities of nuclei between homokaryons and heterokaryons, which had, surprisingly, not been investigated in this model organism yet.

## Material and Methods

### Strains

The information on the strains is given in Supplementary Table S1 (strain ID, species, collection, year, substrate, geographic region of collection, accession number and bioproject number on Genbank). We used 10 heterokaryotic strains of *S. tetrasporum* and 13 heterokaryotic strains of *S. tritetrasporum* stored at -80°C (De Filippo et al., 2025). In the strain ID, the letters A, B, C and D, when present, refer to the spore ID within the isolated tetrads, that were the same as in (De Filippo et al., 2025). We also used 17 heterokaryotic strains of *P. anserina,* from their *MAT1-1* and *MAT1-2* homokaryotic mycelium stored at - 80°C (Silar et al., 2024). Most of these strains were isolated as heterokaryons in 2019-2022 from nature in different areas in France, including the Corsica island, but some were isolated from other countries, including New Zealand, Italy, the Netherlands and Canada. Strains were isolated from soil or from faeces of rabbit, cow, bull, sheep, hare, goat, wallaby or horse. These strains had been stored at -80°C upon isolation. In addition, we used two *S. tetrasporum* strains from the Westerdijk Fungal Biodiversity Institute Collection and the reference *P. anserina* S strain. These three strains may have experienced culture in laboratories, including as homokaryons.

For *S. tetrasporum* and *S. tritetrasporum*, we obtained homokaryotic mycelium of the *MAT1-1* and *MAT1-2* mating types for each heterokaryotic strain stored at -80°C as described previously (Vittorelli et al., 2023, Silar, 2020). Briefly, a plug of heterokaryotic mycelium (5 mm³) was grown for each strain on a V8 medium Petri dish for two to three weeks until perithecia formation. Then, agar Petri dishes were added on the top, upside down, to collect projected spores (Silar, 2020). Homokaryotic spores were then isolated, identified by their smaller size compared to heterokaryotic spores, and grown on Petri dishes with G medium (Silar, 2020) (Fig 1C). These Petri dishes were subjected to a heat shock (30 minutes at 65°C) to induce spore germination. After a few days of growth, mycelia were transferred on M2 medium for growth. The mating-type allele of homokaryotic mycelium grown from those spores was identified by PCR. For this goal, DNA was extracted using Chelex and the mating-type allele (*MAT1-1* or *MAT1-2*) was determined using the PCR conditions described previously (Vittorelli et al., 2023) and the following primer pairs: *MAT1-2* 5’‘GAGGGTGCGAGTCGTGAT’3’ and 3’‘CCAGCCACACGAAAATCC’5’; *MAT1-1* 5’‘GATCGGATTCGCTCCAGA’3’ and 3’‘CGACCGTTGGAAATGACC’5’.

### Genome sequencing and analysis

We used the genomes of 13 of our heterokaryotic strains (CBS815.71, PSN779, PSN881A and PSN710A for *S. tetrasporum,* and PSN352A, PSN571A, PSN1070A, PSN667A, PSN777A, PSN829C, PSN1040, PSN1236 and PSN1433A for *S. tritetrasporum*) available from previous studies (Vittorelli et al., 2023, De Filippo et al., 2025). Genomes have been downloaded at NCBI Genbank under Bioproject numbers PRJNA882797 and PRJNA1190690 (accession numbers in Table S1). DNA had been extracted with the Nucleospin soil kit from Macherey Nagel. Library preparation and genome sequencing had been performed in six different batches by Genewiz, Genoscope or Novogene. The genomes had been sequenced with the Illumina technology (2x150 bp paired-end Novaseq) and had an average coverage of 89X. SNPs were called in (De Filippo et al., 2025). To summarise briefly, we checked Illumina raw read quality using FastQC v0.11.1 v.0.11.5 (Andrews, 2010). Reads were trimmed with Trimmomatic v0.36 (Bolger et al., 2014) in paired-end mode with TruSeq3 adapter and the following options: PE -threads 2 -phred33 TruSeq3-PE.fa:2:30:10 LEADING:10 TRAILING:10 SLIDINGWINDOW:5:10 MINLEN:50.

Illumina trimmed reads were mapped against the CBS815.71-sp3 (MAT1-2) long-read assembly available from the Genbank BioProject accession number PRJNA882797 (assembly accession number JAQKAE000000000 (Vittorelli et al., 2023)) with bowtie2 v2.3.4.1 (Langmead & Salzberg, 2012) with the options -very-sensitive-local -phred33 - X 1000. We used the MarkDuplicates tool from PicardCommandLine v2.8.1 to get rid of duplicated sequences. We called SNPs against the CBS815.71-sp3 assembly with the HaplotypeCaller tool of the Genome Analysis Toolkit (GATK) v4.1.2.0 (McKenna et al., 2010) in the diploid mode. We used CombineGVCF, GenotypeGVCF and SelectVariants to merge data from all strains and conjointly call SNPs. GATK Good Practice rules were followed with the following options: QD=20.0; FS=60.0; MQ=20.0; MQRankSumNeg=-2.0; MQRankSumPos=2.0; <SOR=3.0; QUAL=100. SelectsVariants and VCFtools v0.1.17 were used to filter out indels, and to keep only good-quality SNPs with the 90% missingness option. We predicted SNP effect with SNPeff (Cingolani et al., 2012), recording whether they were synonymous, missense or inducing the gain/loss of stop or start codons, compared to the CBS815.71-sp3 reference genome. We considered that autosomes carried heterozygous regions when at least four consecutive genes carried at least one heterozygous SNP, with less than 50kb between the midpoints of two consecutive genes.

The divergence time between haplotypes in the non-recombining region of *S. tetrasporum*, i.e. a proxy for the age of recombination suppression, was estimated using computed per-gene d_S_ values (synonymous divergence) from a previous study on the *S. tetrasporum* reference genome CBS815.71 (Vittorelli et al., 2023). The number of generations T_g_ since recombination suppression can be estimated by T_g_ = d_S_/2μ, μ being the synonymous substitution rate per generation. Previous estimates of synonymous substitution rates in fungal ascomycetes ranged from 0.9.10^−9^ to 16.7.10^−9^ substitutions per site per year (Kasuga et al., 2002). We used these two values for our computations to obtain a range of dates, as the substitution rate is unknown in *S. tetrasporum*. We considered that *S. tetrasporum* undergoes five generations per year. Indeed, a maximum of 15 generations/year can be obtained in optimal laboratory conditions, which are generally harsher in the wild, with often too low temperature in autumn, winter and spring for growth. To obtain T_y_, the time since recombination suppression in years, we thus use the formula T_y_ = T_g_/5.

### Inoculum preparation for experiments

Prior to each experiment, *MAT1-1* and *MAT1-2* homokaryotic mycelia of the studied heterokaryotic strains were grown on M2 medium (Silar, 2020) for about 7 to 15 days at 27°C, 50% humidity and light and then stored at 4°C for a few days until the beginning of the experiments. On the day of inoculation, fresh inoculum was prepared. Mycelium square plugs of the two mating types for each strain were crushed separately (to form homokaryotic mycelium solutions) and also together (to generate heterokaryotic mycelium solutions; (Silar, 2020, Vittorelli et al., 2023)), with 200 µL of sterile water for *S. tritetrasporum* and *S. tetrasporum* and 150 µL of sterile water for *P. anserina*, using a TissueLyser at 30 Hz for 20 to 30 min.

### Growth speed

Spots of 20 µL of the obtained solutions for homokaryotic mycelium *MAT1-1*, homokaryotic mycelium *MAT1-2* or heterokaryotic mycelium were deposited on Petri dishes with M2 medium (Silar, 2020), with three replicates per strain and condition. One central spot was deposited per Petri dish, corresponding to a replicate. For *S. tritetrasporum*, two batches were performed at different times. Plates were incubated at the conditions of temperature and light exposure detailed in Table 1. Pictures of the plates were taken on two days (see details of the days for each species in Table 1). As the mycelium limits of *P. anserina* were not always clearly distinguishable on pictures, the contour of the mycelial colony was drawn with a marker pen under the Petri dish before taking the picture. Diameters of mycelia were measured using ImageJ 1.54g (Schneider et al., 2012), and the growth speed was computed as the difference in diameter between the two days divided by the number of growth days. We measured one diameter per Petri dish, at the same location for the different time points, using as a reference the label that had been put under the Petri dishes before growth. Some Petri dishes were excluded from the analysis when contamination, dryness or satellite colonies occurred (see Table S2 with raw data).

**Table 1:** Number of strains from *Schizothecium tetrasporum, Schizothecium tritetrasporum* and *Podospora anserina* used for experiments, summary of the conditions (light and temperature) and measure time points for which fitness proxies (growth speed or germination dynamics) were assessed according to species and strains. Information on strains can be found in Table S1.

| Growth speed |  |  |  |  |
| --- | --- | --- | --- | --- |
| Species | Number of strains | Measure time points (days) | Temperature | Light exposure |
| *S. tetrasporum* | 10 | 6 / 14 | 22°C | Yes |
| *S. tetrasporum* | 10 | 6 / 14 | 22°C | No |
| *S. tritetrasporum* | 13 | 6 / 14 | 22°C | Yes |
| *P. anserina* | 17 | 2 / 6 | 18°C | Yes |
| *P. anserina* | 17 | 2 / 6 | 37°C | Yes |
| Germination dynamics |  |  |  |  |
| Species | Number of strains | Measurement time points (days) |  |  |
| *S. tetrasporum* | 8 | 2 / 3 / 7 |  |  |

### Heterokaryotic state persistence and loss of nuclei of one or the other mating type

PCRs were performed on heterokaryotic mycelia 14 days post-inoculation for all *S. tritetrasporum* strains used for measures in the two experiment batches to check whether they remained heterokaryotic. Nine plugs (5 mm³) were sampled in each Petri dish from the inner to the outer of the mycelium ring. DNA was extracted using Chelex and the mating-type allele (*MAT1-1*, *MAT1-2* or both) was determined using the PCR conditions and the primer pairs described above. When only a faint band appeared for one of the two mating types, the PCR was re-run to check the presence of the second band. Some PCRs did not yield any band at all, for neither mating type, and the corresponding plugs were excluded from the analyses (see Table S2 with raw data).

### Number of nuclei per cell in heterokaryons versus homokaryons

We measured the number and density of nuclei per cell in the S strain of *P. anserina* in homokaryons and heterokaryons. Three replicates of each of the *MAT1-1* and the *MAT1-2* homokaryons, as well as of the heterokaryon, were cultured on standard M2 medium for 6 days at 18°C under light. For microscopic analysis, a one centimeter square sample located at the edge of the thallus was cut, fixed in paraformaldehyde (ThermoFischer) 4% in PBT (PBS Phosphate buffered saline + triton X100 (Merck) 0.3%) for 1h, washed in PBT (3X 10 min), stained with 5 µg.mL^-1^ DAPI (4′,6-diamidino-2-phenylindole) in PBT for 30 min then washed in PBT. Samples were mounted inverted on a coverslip in PBT and observed on a Zeiss Axio Observer Z.1 fully motorized inverted microscope; Cool LED pE 400 White; DAPI filter (365 D395 445/50); objective 63x/1.4 Plan-Apo Oil, WD: 0.19mm, ∞/0.17. For each sample, observations were made in the two first millimeters from the edge of the thallus (growth zone). Images were treated with the Fiji software (Schindelin et al., 2012). A total of 30 articles (the hyphal cells delimited by two septa) were selected at random in each replicate, to avoid any biased selection of articles depending on their number of nuclei, and their length was measured in the transmitted light canal. Then, the number of nuclei per article was counted and the density was computed as the number of nuclei divided by the length of articles (see Table S3 with raw data).

### Germination dynamics

For the 10 heterokaryotic strains of *S. tetrasporum*, three spots of 20 µL of heterokaryotic mycelium solution were deposited on a single Petri dish with V8 medium (Vittorelli et al., 2023) and incubated at 22°C until perithecia formation (about 15 days). PSN881A and PSN779 strains did not produce any perithecia and were excluded from the analysis. Agar Petri dishes were added on the top, upside down, to collect projected spores (Vittorelli et al., 2023). We isolated 10 large spores and 10 small spores per heterokaryotic strain, which were deposited on a Petri dish with G medium (Silar, 2020), allowing germination (two batches per strain). Small spores are homokaryotic, while large spores are heterokaryotic at the mating-type locus with a 90% probability, as estimated in (Vittorelli et al., 2023) based on genotyping of the germinated mycelium, which is a minimum estimate, as the mycelium could also have lost a nucleus during growth before genotyping. These Petri dishes were foiled with aluminium paper, subjected to a heat shock of 30 min at 65°C and incubated at 27°C to induce germination (Silar, 2020). Germinated spores were counted on days 2, 3 and 7 to analyse the germination dynamics of small and large spores, i.e., the rates of germination over time (Table 1). A few spores were excluded from the analysis when there was contamination or two spores were observed germinating instead of one (see Table S4 with raw data).

### Graphical summaries and statistical analyses

All statistical analyses and graphical displays were carried out using the R software v 4.3.2 (R Core Team, 2023). A statistical significance level of 5% was considered for all statistical tests. The linearity of the response and predictor relationship and the Gaussian distribution of the residuals in all models (including random effects intercept and slope in the case of random-effects models) were assessed visually using R built-in facilities (fitted vs. residual values plot and QQ-plot, respectively).

For the growth experiment analyses in *S. tritetrasporum*, *S. tetrasporum* and *P. anserina*, linear mixed-effect models were fitted, using the restricted maximum likelihood method from the lme4 package, considering as fixed effects either ploidy (i.e., heterokaryon or homokaryon) or the nuclear status (i.e., heterokaryon, *MAT1-1* homokaryon or *MAT1-2* homokaryon), in addition to the growing condition, when applicable (i.e. light or dark for *S. tetrasporum*, 18°C or 37°C for *P. anserina*). The interaction between the growing condition and the ploidy or the nuclear status was also included in the models. The strain ID was considered a random effect in all models, as well as the batch in the case of *S. tritetrasprorum*. The statistical significance of fixed effects was assessed using F-tests with the lmerTest package and a Satterthwaite approximation for degrees of freedom. The emmeans package was used to estimate marginal means and their associated standard errors and to estimate predefined pairwise contrasts, with degrees of freedom calculated using the Kenward-Roger method. Obtained p-values from Student t-tests were adjusted using the Holm method to account for multiple testing.

For the analysis testing the preferential loss of nuclei of one or the other mating type in heterokaryotic mycelia in *S. tritetrasporum*, a generalized linear mixed-effects model with a Binomial distribution was used. The model was designed to assess the proportion of plugs in which nuclear loss occurred, with the genotype of the plug after the loss (i.e., *MAT1-1* or *MAT1-2*) as a fixed effect and strain ID and batch as random effects. For this analysis, a generalized linear mixed-effect model using the maximum likelihood method with the Laplace approximation for random effects was fitted with the lme4 package. The statistical significance of the fixed effect was assessed using a χ2-test with the car package. The ggeffect package was used to estimate marginal means and their associated standard errors. The emmeans package was used to estimate a predefined pairwise contrast using a z-test.

For nucleus number analysis in the S strain of *P. anserina*, a generalised linear model with a Poisson distribution was used, modelling the number of nuclei per article considering the nuclear status (i.e., heterokaryon, *MAT1-1* homokaryon or *MAT1-2* homokaryon) as a fixed effect. The statistical significance of the fixed effect was assessed using a χ2-test with the car package. The ggeffect package was used to estimate marginal means and their associated standard errors. The emmeans package was used to estimate predefined pairwise contrasts and obtained p-values from z-test were adjusted using the Holm method to account for multiple testing. A proxy of the nucleus density was computed as the number of nuclei divided by the size of the article, and a linear model was fitted, modelling the density of nuclei per article considering the nuclear status (i.e., heterokaryon, *MAT1-1* homokaryon or *MAT1-2* homokaryon) as a fixed effect. The statistical significance of the fixed effect was assessed using a F-test. The emmeans package was used to estimate marginal means and their associated standard errors and to estimate predefined pairwise contrasts. Obtained p-values from Student t-tests were adjusted using the Holm method to account for multiple testing.

For the germination experiment analysis in *S. tetrasporum*, a generalised linear mixed-effect model with a Binomial distribution was used, modelling germination rate as a linear combination of day (as a continuous predictor) and spore size (i.e. large or small), treated as fixed effects, and strain ID and batch as random effects. For this analysis, a generalized linear mixed-effect model using the maximum likelihood method with the Laplace approximation for random effects was fitted with the lme4 package. The statistical significance of fixed effects was assessed using χ2-tests with the car package. The ggeffect package was used to estimate marginal means and their associated standard errors. The emmeans package was used to estimate a predefined pairwise contrast using a z-test.

## Results

### Genome-wide homozygosity except in the mating-type locus region, that carries likely heterozygous deleterious mutations

To assess whether heterozygosity in the genomes of the strains used for experiments was mainly located around the mating-type locus and whether signs of heterozygous deleterious mutations could be detected, we used the available genomes of four and nine heterokaryotic strains of *S. tetrasporum* and *S. tritetrasporum* (De Filippo et al., 2025). All strains displayed heterozygosity around their mating-type locus, in a genomic region ranging from 1.38Mb to 1.47Mb in *S. tetrasporum* and 1.30Mb to 1.71Mb kb in *S. tritetrasporum* (Figs 2, S1 and S2).

**Figure 2:**
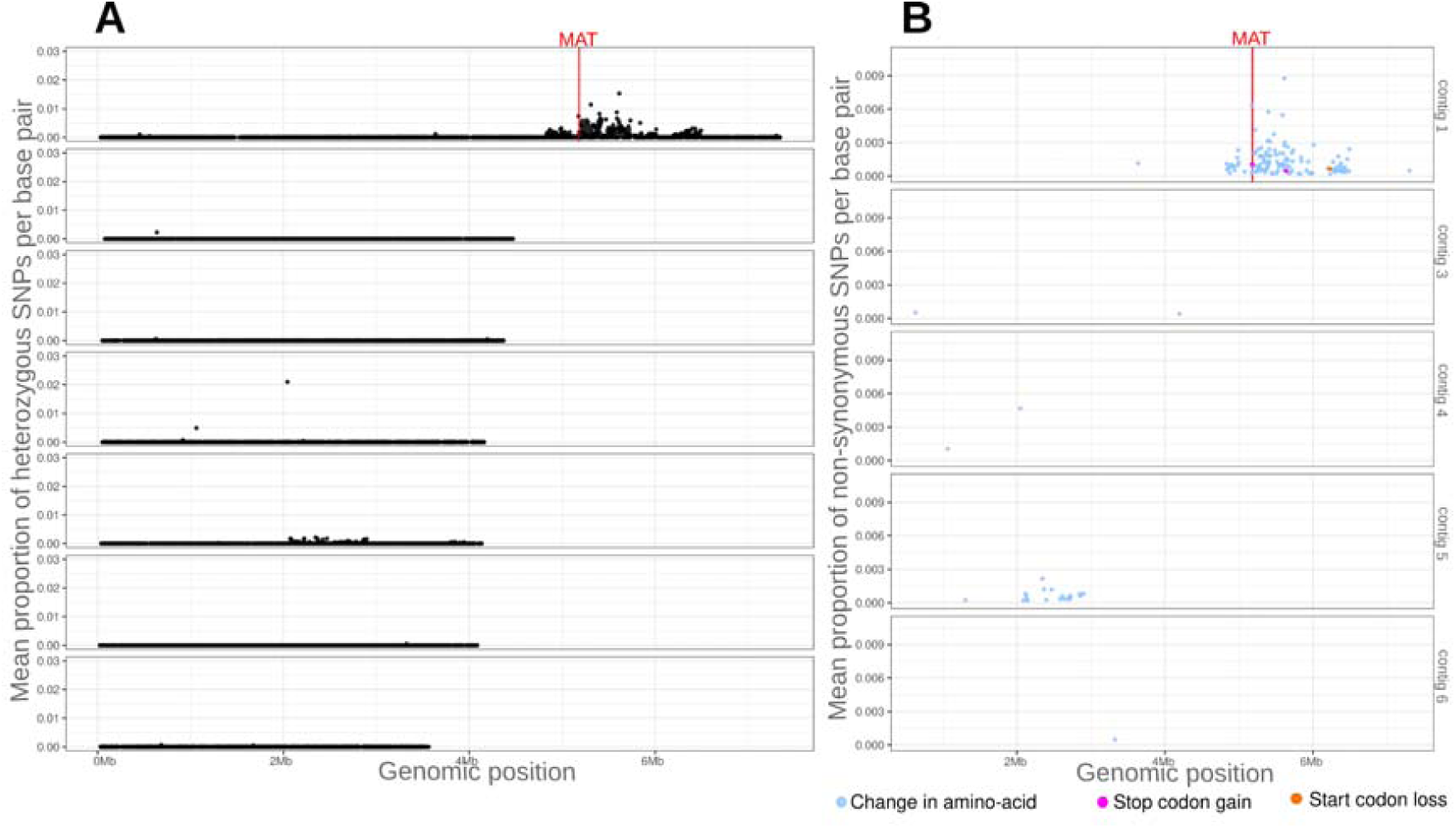
Genomic analyses of heterozygosity and mating-type specific putative deleterious mutations (non-synonymous SNPs) along the chromosomes of the *Schizothecium tritetrasporum* PSN829 strain, as an illustration of the situation with genome-wide homozygosity except a few heterozygous autosomal regions in addition to the flanking region of the mating-type locus. A) Genome-wide per-gene density in heterozygous SNPs. Each dot represents one gene, the Y axis corresponding to the mean proportion of heterozygous SNPs per base pair for each gene, and the X axis to the genomic position along the CBS815.71-sp3 assembly in megabases (Mb). The seven longest contigs in the assembly are represented, by length order. B) Genome-wide per-gene density in heterozygous non-synonymous SNPs. Each dot represents one gene, the Y axis corresponding to the mean proportion of heterozygous non-synonymous SNPs per base pair; these SNPs introduced either a missense mutation (i.e., change in amino-acid; blue dot), the gain of a stop codon (pink dot) or the loss of a start codon (orange dot) compared to the reference genome CBS815.71-sp3. The X axis corresponds to the genomic position along the CBS815.71-sp3 assembly in megabases (Mb). Dots indicating a null value have been removed for clarity. The mating-type locus (MAT) position is symbolized by a vertical red line.

As reported for the CBS815.71 *S. tetrasporum* reference strain (Vittorelli et al., 2023), distinct evolutionary strata seemed to be present in all strains, i.e. segments with different divergence levels (Figs 2, S1 and S2). We estimated the age of recombination suppression in *S. tetrasporum* based on previous estimates of synonymous differences in the non-recombining region in the reference genome CBS815.71 (Vittorelli et al., 2023, Hartmann et al., 2021a) and synonymous substitutions rates in ascomycetes (Kasuga et al., 2002), to be between 40 ky and 730 ky for the oldest stratum, and between 9 ky and 170 ky for the youngest stratum.

Among the genomes of the 13 heterokaryotic strains analysed, only four showed some tracks of heterozygosity in autosomes, corresponding to one *S. tetrasporum* strain (PSN710A; Fig S1) and three *S. tritetrasporum* strains (PSN777A, PSN352A and PSN829C; Figs 2 and S2); the heterozygous regions were short: on average, the strains with heterozygosity on the autosomes had a total of 910 kb of heterozygous regions on their autosomes, i.e., only 3.6% of the total autosomal genomic size. The nine other strains were homozygous genome-wide, except in the non-recombining region near the mating-type locus (Figs S1 and S2).

For all strains analysed, the region around the mating-type locus also harbored heterozygous non-synonymous mutations, especially changes in amino-acids, but also gains of stop codons and losses of start codons (Figs 2, Fig S1 and Fig S2), suggesting that deleterious mutations may be present at the heterozygous state. We detected no heterozygous SNP inducing the loss of a stop codon or the gain of a start codon compared to the reference genome.

### Growth advantage of heterokaryons

To investigate the presence of sheltered load in the non-recombining regions around the mating-type locus, or at their margin, in *S. tetrasporum*, *S. tritetrasporum* and *P. anserina,* experiments were performed to compare the fitness of heterokaryons to that of homokaryons using strains as biological replicates in each species (10, 13 and 17 strains, respectively; Table S1; Fig 1).

For *S. tetrasporum* and *S. tritetrasporum*, we used as fitness proxies the mycelium growth speed under light at 22°C between day 6 and day 14. In addition, we used as fitness proxies for *S. tetrasporum* the mycelium growth speed under dark at 22°C and assessed the spore germination dynamics between day 2 and day 7. For *P. anserina,* we measured mycelium growth speed between day 2 and day 6 under light at 18°C and 37°C. These conditions were chosen for each species based on preliminary experiments or previous knowledge on the different species, to use more or less stressful conditions. Table S2 gives the raw measures for the growth experiment for which 8% of plates have been excluded on average per species due to the presence of satellite colonies, contaminants or dryness.

In *S. tritetrasporum*, heterokaryons grew significantly faster than homokaryons under light at 22°C, by 3.7% (see Fig 3A showing raw data, Fig 3B showing model estimates, Table S5A). When considering the two types of homokaryons separately in the statistical analysis, we found that heterokaryons grew significantly faster than *MAT1-1* homokaryons, by 5.7%. *MAT1-2* homokaryons had a tendency to grow more slowly than heterokaryons, but the difference was not significant (Fig 3A showing raw data, Fig 3C showing model estimates, Table S5B).

**Figure 3:**
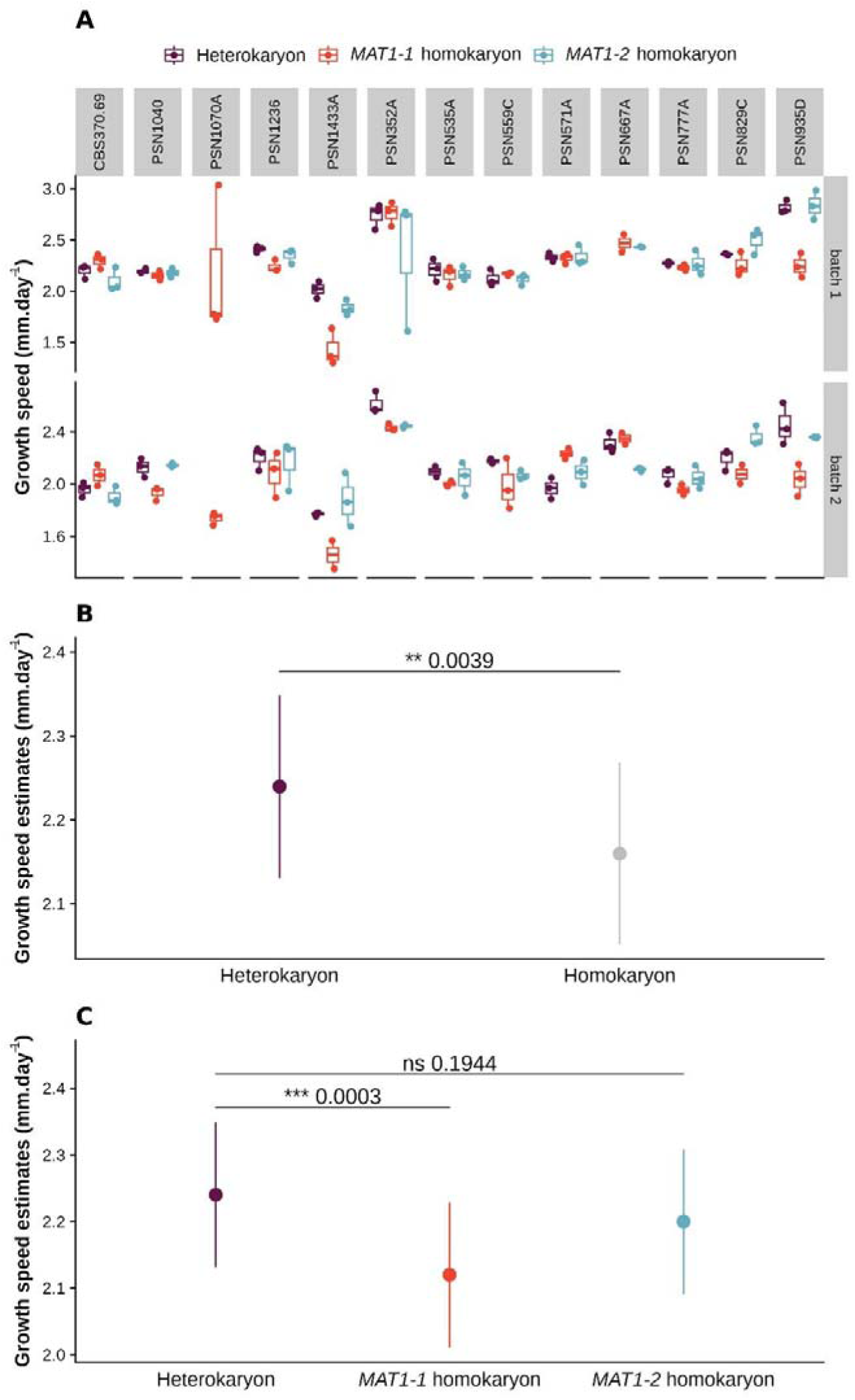
Mycelium growth speed of heterokaryons versus homokaryons in *Schizothecium tritetrasporum.* **A)** Mycelium growth speed (mm.dayL¹) of heterokaryons (in purple), *MAT1-1* homokaryons (in red) and *MAT1-2* homokaryons (in blue), of the three replicates for each of two batches, for the 13 strains of *S. tritetrasporum,* between day 6 and day 14, grown under light at 22°C. Boxplot elements: central line: median, box limits: 25th and 75th percentiles, whiskers: 1.5× interquartile range. **B)** Model estimates of mycelium growth speed for heterokaryons (in purple) and homokaryons (in grey) in *S. tritetrasporum.* Dots and bars represent marginal means and standard errors estimated from a linear mixed-effect model explaining growth speed using ploidy (i.e. heterokaryon or homokaryon) as a fixed effect, and strain ID and batch as random effects. **C)** Model estimates of mycelium growth speed for heterokaryons (in purple), *MAT1-1* homokaryons (in red) and *MAT1-2* homokaryons (in blue) in *S. tritetrasporum.* Dots and bars represent marginal means and standard errors estimated from a linear mixed-effect model explaining growth speed using the nuclear status (i.e. *MAT1-1* homokaryon, *MAT1-2* homokaryon or heterokaryon) as a fixed effect, and strain ID and batch as random effects. P-values correspond to pairwise t-tests adjusted for multiple testing using the Holm method (ns: non-significant; **: p<0.01; ***: p<0.001).

In *S. tetrasporum*, heterokaryons also grew significantly faster than homokaryons under light at 22°C, by 3.6% (see Fig 4A showing raw data, Fig 4B showing model estimates, Table S5C; the global model was not significant, but individual tests for pairwise comparisons, relying on unpooled variance between all conditions, indicate a significant difference between homokaryons and heterokaryons for growth speed under light). The difference in growth speed between homokaryons and heterokaryons was not significant under dark conditions, although the tendency was similar (Fig 4A and B, Table S5C). When separating the two homokaryons in the analysis, we found that heterokaryons grew significantly faster than *MAT1-1* homokaryons under light at 22°C, by 5.3% (see Fig 4A showing raw data, Fig 4C showing model estimates, Table S5D). Heterokaryons had a tendency to also grow faster than MAT1-2 homokaryons under both light and dark conditions, albeit the difference was not significant (Figs 4A and C, Table S5D).

**Figure 4:**
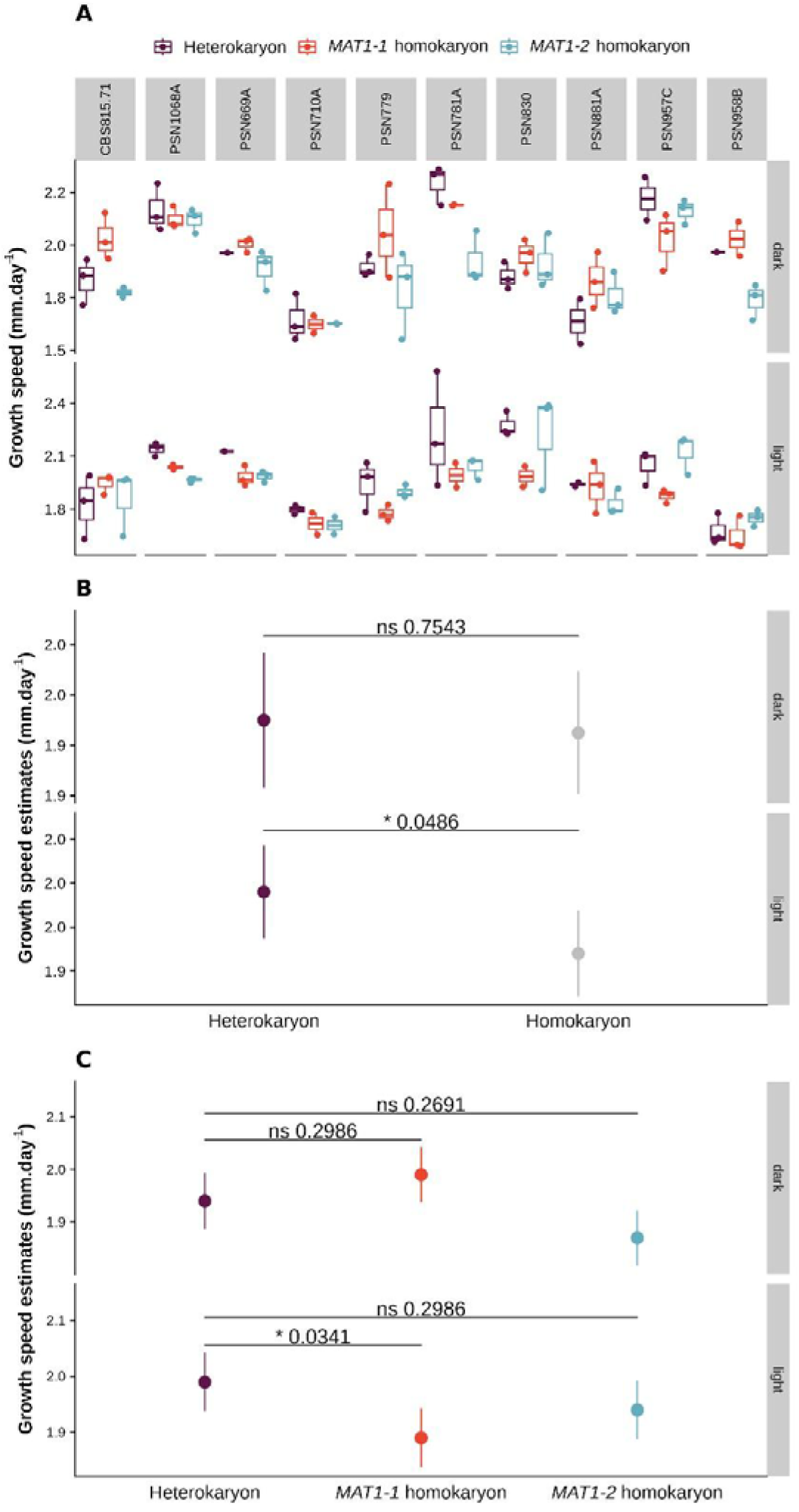
Mycelium growth speed of heterokaryons versus homokaryons in *Schizothecium tetrasporum*. **A)** Mycelium growth speed (mm.dayL¹) of heterokaryons (in purple), *MAT1-1* homokaryons (in red) and *MAT1-2* homokaryons (in blue), of the three replicates of the 10 *S. tetrasporum* strains between day 6 and day 14, grown under light or dark conditions at 22°C. Boxplot elements: central line: median, box limits: 25th and 75th percentiles, whiskers: 1.5× interquartile range. **B)** Model estimates of mycelium growth speed for heterokaryons (in purple) and homokaryons (in grey) in *S. tetrasporum* under light or dark conditions. Dots and bars represent marginal means and standard errors estimated from a linear mixed-effect model explaining growth speed using ploidy (i.e. homokaryon or heterokaryon) and condition (i.e. light or dark) as fixed effects, with their interaction, and strain ID as a random effect. **C)** Model estimates of mycelium growth speed for heterokaryons (in purple), *MAT1-1* homokaryons (in red) and *MAT1-2* homokaryons (in blue) in *S. tetrasporum* under light or dark conditions. Dots and bars represent marginal means and standard errors estimated from a linear mixed-effect model explaining growth speed using nuclear status (i.e. *MAT1-1* homokaryon, *MAT1-2* homokaryon or heterokaryon) and condition (i.e. light or dark) as fixed effects, with their interaction, and strain ID as a random effect. P-values correspond to pairwise t-tests adjusted for multiple testing using the Holm method (ns: non-significant; *: p<0.05).

In *P. anserina*, heterokaryons had a tendency to grow faster than homokaryons at the two temperatures, but no differences were significant when considering both types of homokaryons together (see Fig 5A showing raw data, Fig 5B showing model estimates, Table S5E). However, when considering the two types of homokaryons separately, heterokaryons grew significantly faster than *MAT1-2* homokaryons at 18°C, by 7.9% (see Fig 5A showing raw data, Fig 5C showing model estimates, Table S5F). The tendency was similar at 37°C but not significantly so (Figs 5A and 5C, Table S5F).

**Figure 5:**
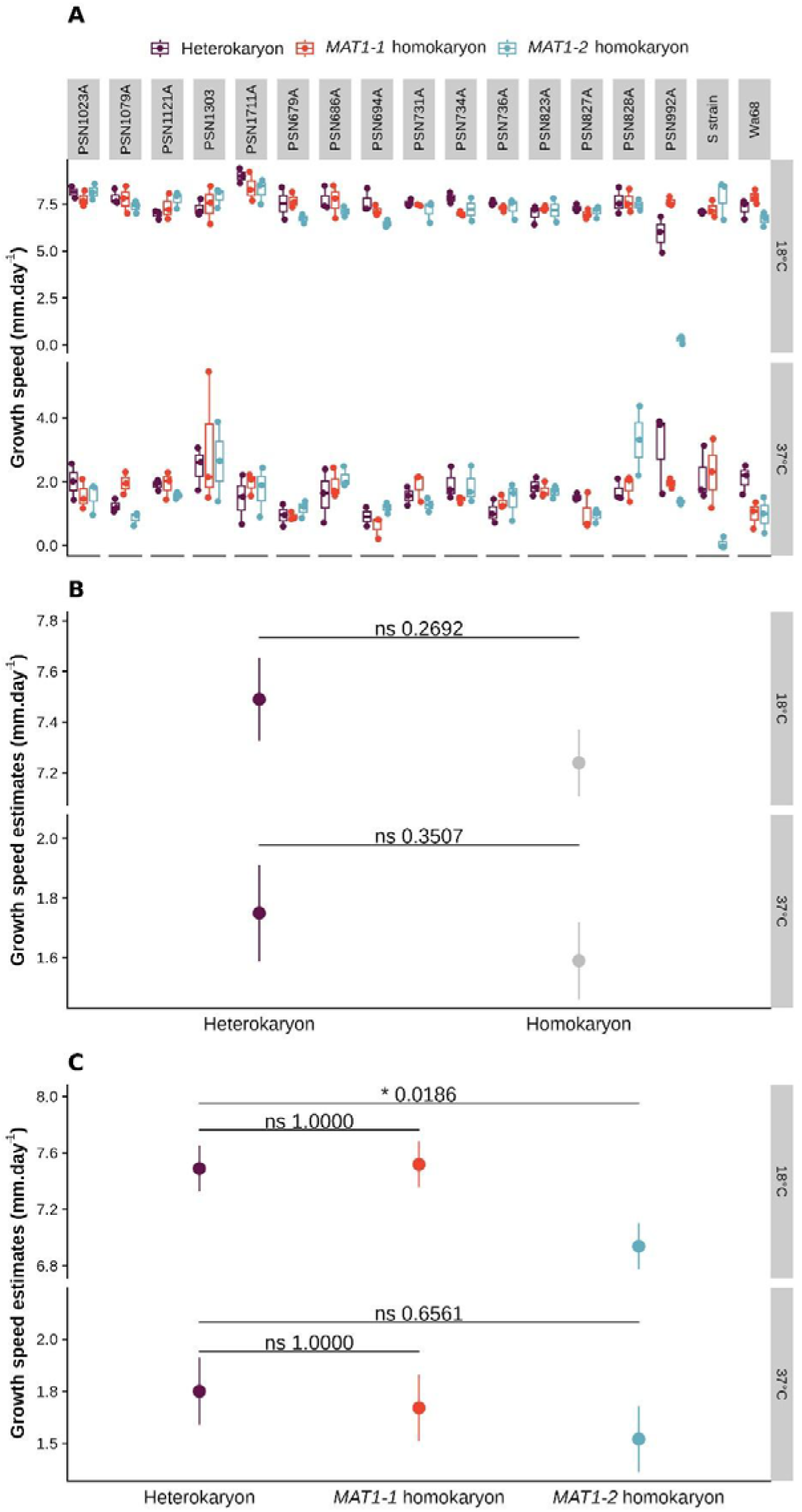
Mycelium growth speed of heterokaryons versus homokaryons in *Podospora anserina*. **A)** Mycelium growth speed (mm.dayL¹) of heterokaryons (in purple), *MAT1-1* homokaryons (in red) and *MAT1-2* homokaryons (in blue), of the three replicates of the 17 *P. anserina* strains between day 2 and day 6, grown at 18°C or 37°C under light conditions. Boxplot elements: central line: median, box limits: 25th and 75th percentiles, whiskers: 1.5× interquartile range. **B)** Model estimates of mycelium growth speed for heterokaryons (in purple) and homokaryons (in grey) in *P. anserina* at 18°C and 37°C. Dots and bars represent marginal means and standard error estimated from a linear mixed-effect model explaining growth speed using ploidy (i.e. homokaryon or heterokaryon) and condition (i.e. 18°C or 37°C) as fixed effects, with their interaction, and strain ID as a random effect. **C)** Model estimates of mycelium growth speed of heterokaryons (in purple), *MAT1-1* homokaryons (in red) and *MAT1-2* homokaryons (in blue) in *P. anserina* at 18°C and 37°C. Dots and bars represent marginal means and standard errors estimated from a linear mixed-effect model explaining growth speed using nuclear status (i.e. *MAT1-1* homokaryon, *MAT1-2* homokaryon or heterokaryon) and condition (i.e. 18°C or 37°C) as fixed effects, with their interaction, and strain ID as a random effect. P-values correspond to pairwise t-tests adjusted for multiple testing using the Holm method (ns: non-significant; *: p<0.05).

Re-running all statistical analyses after removing the strains having likely experienced laboratory culture (i.e., CBS370.69, CBS815.71 and the S strain) and those showing traces of heterozygosity in autosomes (i.e. PSN710A, PSN777A, PSN352A and PSN829C) did not change the conclusions: the same fixed effects and pairwise differences as above remained significant (not shown). In *S. tetrasporum,* this even led to a significant effect of ploidy on mycelium growth, and rendered the global model significant.

### Heterokaryotic mycelia lost nuclei of one mating type in small proportions, and with no preferential loss for one mating type over the other

Given that some level of nuclear loss of one mating type has been reported to occur during heterokaryotic mycelium growth in the *S. tetrasporum* species complex (Vittorelli et al., 2023), while it does not occur in *P. anserina*, we investigated whether, and to what extent, such a phenomenon occurred in the *S. tritetrasporum* heterokaryotic mycelia used for growth experiments. For this goal, we genotyped the mating-type locus for nine plugs per Petri dish. Table S2 gives the PCR results, 3% of the plugs from plates of *S. tritetrasporum* heterokaryotic mycelium used for growth measures having been excluded because PCRs did not yield any band at all, for neither mating type.

We found that mycelia had remained mostly heterokaryotic, although we detected a single mating type in 27.7% of the plugs (Figs S3 and S4A). In these plugs where a single mating-type was detected, there was no significant over-representation of one mating type over the other one (see Fig S4A showing raw data, Fig S4B showing model estimates, Table S5G).

### No differences in nuclei number or density between heterokaryotic and homokaryotic mycelia

As nucleus number per cell might vary between heterokaryotic and homokaryotic mycelia, then possibly impacting their growth rates, we investigated whether the number or the density of nuclei per cell differed between heterokaryons and each of the homokaryons in the reference *P. anserina* S strain. Table S3 gives the raw measures of nucleus counts and article lengths, articles being hyphal cells delimited by two septa.

The differences in the number and density of nuclei per article between heterokaryons and each of the homokaryons in the *P. anserina* S strain were small and not significant (Fig S5 showing a picture of nucleus counts; Figs S6A and S6C showing raw data, Figs S6B and S6D showing model estimates; Tables S5H and S5I).

### No differences in germination rates between heterokaryotic and homokaryotic spores

We also investigated the existence of a heterokaryon fitness advantage in terms of spore germination, using the same *S. tetrasporum* strains as for growth experiments. Table S4 gives the raw measures for the germination experiment, for which two out of the 10 strains of *S. tetrasporum* were excluded due to lack of perithecia and less than 1% of the spores were excluded, due to contamination or the germination of two spores, meaning that two spores had been deposited instead of one.

The comparison of the germination dynamics of small spores (i.e. homokaryotic spores) versus large spores (i.e. mainly heterokaryotic spores; (Vittorelli et al., 2023)) in *S. tetrasporum* did not reveal any evidence of heterozygous advantage. Indeed, although large spores had a tendency to germinate at a higher rate than small spores, the difference in germination rate between small and large spores was not significant (see Fig 6A showing raw data, Fig 6B showing model predictions, Table S5J). Adding an interaction between the two fixed effects, i.e. spore size and day, and/or considering the variable day as a factor in the statistical model did not alter the previous conclusions (not shown).

**Figure 6:**
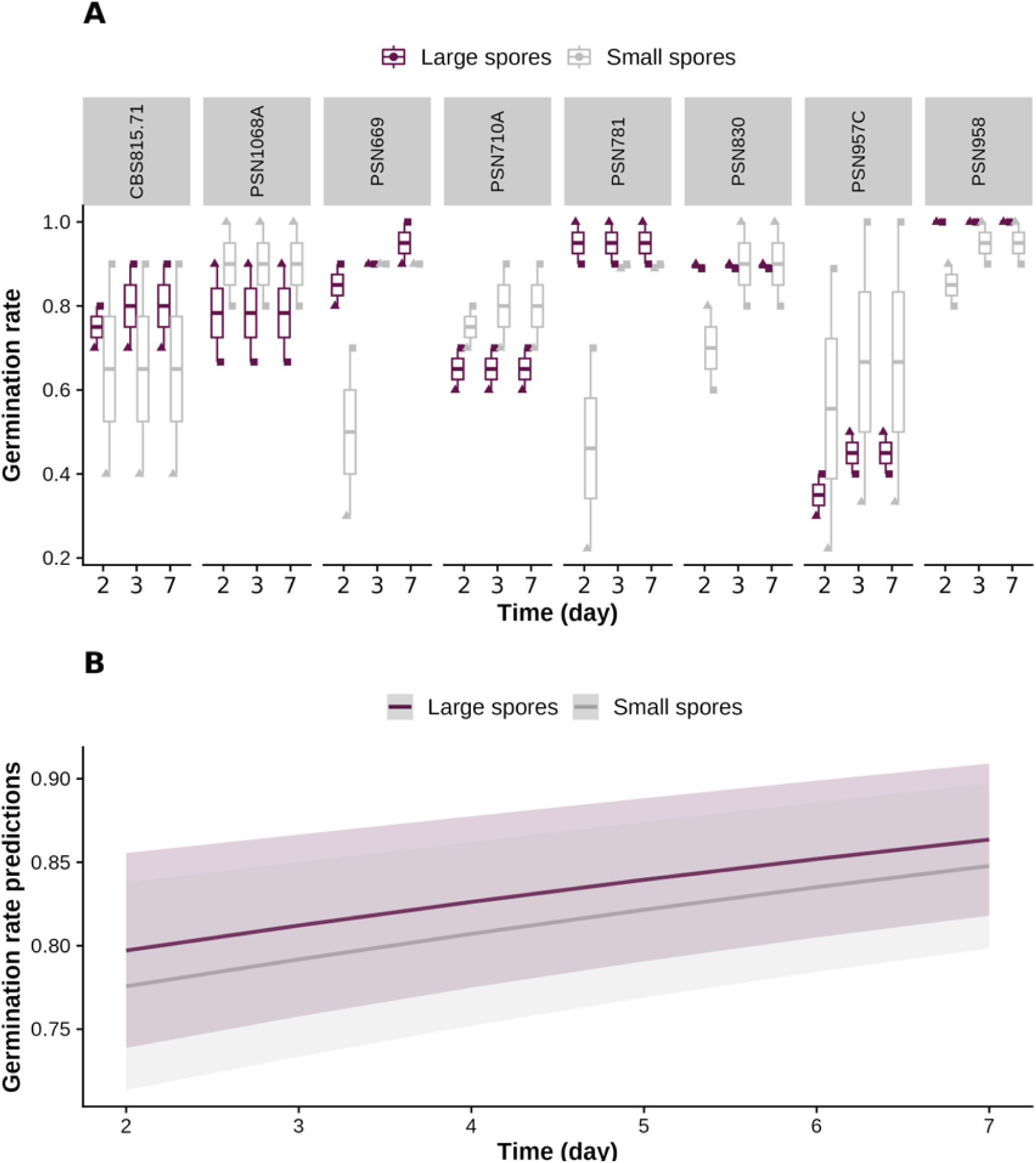
Spore germination dynamics in *Schizothecium tetrasporum,* depending on their heterokaryotic versus homokaryotic nuclear status, as inferred from their size. **A)** Germination rate on days 2, 3 and 7 for small spores (in purple) and large spores (in grey) in the 10 spores for each of two batches of the eight *S. tetrasporum* strains. Symbol shapes correspond to batches. Boxplot elements: central line: median, box limits: 25th and 75th percentiles, whiskers: 1.5× interquartile range. **B)** Model predictions of germination rate dynamics for small spores (in purple) and large spores (in grey) in *S. tetrasporum*. Curves and shaded error bands represent model prediction and standard errors from the fit of the germination dynamics using a generalised linear mixed-effect model with a Binomial distribution modelling germination rate as a linear combination of day (as a continuous predictor) and spore size (i.e. large or small) treated as fixed effects, and strain ID and batch as random effects (pairwise z-test between the two spore size levels: p=0.4459, non-significant).

## Discussion

### Likely presence of recessive deleterious mutations around the mating-type locus

Our fitness assay showed that heterokaryons were fitter than at least one homokaryon under light at 18°C in *P. anserina* and at 22°C in *S. tetrasporum* and *S. tritetrasporum*, suggesting the presence of recessive deleterious mutations in the non-recombining region around the mating-type locus, and/or tightly linked to it, being associated to a specific single mating-type allele. We indeed confirmed that *S. tetrasporum* and *S. tritetrasporum* strains were mostly homozygous genome-wide, except around the mating-type locus, as previously shown in *P. anserina* (Hartmann et al., 2021a). We further revealed the existence of heterozygous non-synonymous mutations in the non-recombining region, in particular changes in amino-acids, gains of stop codons and losses of start codons, that constitute potential deleterious mutations. Our findings thus indicate the existence of recessive deleterious mutations around the mating-type locus, that are expressed at the homokaryotic state, but sheltered at the heterokaryotic state, and that impact more homokaryons of a mating type than the alternative mating type, and only under particular conditions. Note that, even if the sheltered load was only detected at some temperature, under light and during growth (and not germination), it still means that sheltered load exists. The sheltered load caused 3-8% of decrease in mycelium growth speed, which represents a non-negligible fitness impact, especially for colonizing a dung before competitors or before it dries out. Our study is one of the rare experiments showing the existence of a sheltered load in sex-related chromosomes. Only a few studies could investigate a sheltered load so far, around self-incompatibility loci in plants (Stone, 2004, Llaurens et al., 2009, Le Veve et al., 2024) as this is one of the rare systems where mating-type loci can be rendered homozygous. To investigate the presence of a sheltered load, we leveraged another unique biological system, pseudo-homothallic fungi, whose life cycle allows fitness comparison between individuals in which mutations around the mating-type locus are sheltered versus exposed to selection, while the rest of the genome is exposed to selection, being highly homozygous due to high selfing rates in nature. Previous evidence of a sheltered load in fungal-like organisms include haplo-lethal alleles linked to mating types in anther-smut fungi and oomycetes, in which one or the two mating types fail to grow at the haploid stage or cause a segregation bias in progenies (Hood & Antonovics, 2000, Thomas et al., 2003, Oudemans et al., 1998, Judelson et al., 1995). The non-recombining regions were, however, much older in these cases, i.e., million years old (Duhamel et al., 2022, Duhamel et al., 2023), while we estimated here that recombination suppression occurred in *S. tetrasporum* ca. 730 ky ago. Few other studies have compared fitness proxies between homokaryons and heterokaryons in fungi. In three sibling species of the species complex *Neurospora tetrasperma*, other Sordariales fungi with recombination suppression around its mating-type locus, the quantity of asexual haploid spores (conidia) produced by mycelia has been compared between heterokaryons and homokaryons (Meunier et al., 2018). No evidence of sheltered load was found, but a single strain has been used for these experiments in each of the sibling species (Meunier et al., 2018). Recessive deleterious mutations have also been experimentally detected in a few autosomal supergenes (Jay et al., 2021, Wielstra, 2020, Hill et al., 2023). Altogether, such direct experimental evidence of sheltered load is important for understanding the evolution of non-recombining regions.

### Sheltered load detected in homokaryons of a single mating type and not in all conditions

The lower fitness of homokaryons was not detected in all traits or all conditions, and was significant in a single one of the homokaryons, in all three species tested. This may be explained by the young age of the recombination suppression, having allowed the accumulation of only few deleterious mutations so far, in a few genes, perhaps only involved in tolerance to particular conditions, and by chance more in the flanking region of one of the mating-type alleles. The genomic analyses in fact suggest differences between mating-type chromosomes in terms of non-synonymous substitutions.

A possible non-negligible haploid phase can also purge, to some extent, recessive mutations with large detrimental effects on some traits and in some specific conditions. Indeed, although rare and with an unknown importance in nature, some homokaryotic spores are produced in these fungi: in *S. tetrasporum*, 2% to 10% of asci with small (i.e. homokaryotic) spores have been reported (Raju & Perkins, 1994, Vittorelli et al., 2023); in *P. anserina*, 5% of spores are homokaryotic (Ament-Velásquez & Vogan, 2022). The loss of nuclei of one mating type during mycelium growth, as shown here and previously (Vittorelli et al., 2023), may also expose recessive deleterious mutations to selection.

The growth as haploids could be linked to laboratory conditions, as a few of the strains used in our experiments have been maintained in laboratories for some time (in particular the *P. anserina* S strain), where they can be cultivated as homokaryons. If these strains have experienced purifying selection as homokaryons more often than wild strains, this may have reduced fitness differences between homokaryons and heterokaryons in some conditions; the effect would therefore be conservative regarding the differences that we detected. In addition, stock strains are maintained at -80°C, from where they are taken for each experiment, likely limiting this effect. Furthermore, removing these strains from analyses did not change our results.

Another mechanism allowing the purge of deleterious mutations could be the occurrence of very rare events of recombination in the MAT-proximal region, as reported in *P. anserina* (Contamine et al., 1996). Such extremely rare events of recombination, also reported in sex chromosomes in frogs (Rodrigues et al., 2018), has been proposed to indeed prevent strong differentiation and high level of degeneration of sex chromosomes (Rodrigues et al., 2018).

An explanation for a stronger sheltered load on homokaryons of one mating type could be an over-representation of nuclei of one of the two mating types in heterokaryons, exposing more the genomic background of this mating type to selection. Similarly, if the genome of one of the two nuclei in each cell, associated to a particular mating type, is expressed at a higher level, as suggested in *N. tetrasperma* (Meunier et al., 2022), selection could act more strongly on this mating-type’s genomic background. The effects of higher accumulation of deleterious mutations by chance in one mating type’s background and of stronger selection on the other mating type’s background could be mutually reinforcing: the first partly recessive deleterious mutations occurring by chance more in the non-recombining regions of one mating type may make nuclei of this mating type less expressed or replicating more slowly within cells, being then under-represented, leading to stronger selection on the genomic background of the other mating-type.

Our experiments actually suggest that nuclear losses of one mating type occur in sectors of heterokaryotic mycelia in *S. tritetrasporum*, as previously reported in the *S. tetrasporum* species complex (Vittorelli et al., 2023), but none of the mating types showed a preferential tendency to be lost. Such occasional losses of nuclei mean that some parts of the mycelium in the heterokaryotic treatment may actually be homokaryotic. Such a phenomenon, if any effect in our experimental design, would have attenuated differences between homokaryons and heterokaryons, thereby possibly preventing the detection of lower homokaryon fitness in some conditions or making their disadvantage appear weaker than it is. In addition, large spores have been shown to be sometimes homokaryotic in *S. tetrasporum* (less than 13% of large spores germinated as homokaryons, which may also be due to nucleus loss during growth (Vittorelli et al., 2023)); this could have artefactually reduced the difference between homokaryotic and heterokaryotic spores and have prevented the detection of a lower fitness of homokaryons for spore germination dynamics.

### Alternative hypotheses to sheltered load

There may be, in principle, alternative hypotheses to sheltered load explaining the lower fitness of homokaryons. Although the species studied mainly reproduce by selfing and most strains display genome-wide homozygosity (Grognet et al., 2014, Hartmann et al., 2021a, Vittorelli et al., 2023), some rare events of outcrossing can generate short tracks of heterozygosity in genomic regions far from the mating-type locus. The analysis here of the genomes of 13 heterokaryotic strains however showed that most strains displayed genome-wide homozygosity, except around the mating-type locus, in *S. tetrasporum* and *S. tritetrasporum*. Removing the strains with short tracks of heterozygosity in autosomes and the laboratory strains from statistical analyses did not affect the general results, indicating that the lower fitness of homokaryons was not due to these autosomal heterozygous regions. Similar patterns have previously been reported in *P. anserina*: only two out of ten strains had small heterozygous regions in autosomes (Hartmann et al., 2021a). The panel of *P. anserina* strains analysed in the previous study only had the S strain in common with the set of strains used here, and the S strain displays genome-wide homozygosity except around its mating-type locus (Grognet et al., 2014).

Other types of heterozygous advantage can also exist, although such advantages are rare in nature and mostly related to host-parasite interactions (Gemmell & Slate, 2006), and therefore unlikely in our experimental setting. The lower fitness of homokaryons could also have been due, in principle, to a ploidy advantage for heterokaryons, for example related to gene expression dosage, which may have been expected to be twice as high in heterokaryons as in homokaryons. We, however, found no evidence for a higher number or higher density of nuclei per cell in heterokaryons compared to homokaryons in the S strain of *P. anserina,* which had never been investigated so far in this model organism. Furthermore, this hypothesis would predict a lower fitness of both homokaryons compared to heterokaryons, while we found significance only between one homokaryon and the heterokaryon.

Another explanation for differences in growth between heterokaryons and homokaryons could be differences in their developmental programs. No data or observation are available to our knowledge to support this hypothesis, however, and similar expression patterns have been found in homokaryons and heterokaryons in *N. tetrasperma*, except in the non-recombining region (Meunier et al., 2022). The gene expression differences in *N. tetrasperma* between homokaryons and heterokaryons (Meunier et al., 2022) are indeed restricted to genes around the mating-type locus, which is likely due to degeneration following less efficient selection (Fontanillas et al., 2015) rather than different developmental programs resulting from selection for different behaviours. Here too, this hypothesis would predict a heterokaryon advantage over each of the homokaryons.

## Conclusion

In conclusion, our findings based on both genome analyses and experiments suggest the presence of a sheltered load, i.e., of recessive deleterious mutations at the heterozygous state around the mating-type locus, associated to a single mating type, in three species of Sordariales fungi with young regions of recombination suppression. Leveraging the experimental assets of fungi, allowing cultivating separately haploid-like and diploid-like life stages, in a system with two mating types permanently heterozygous and genome-wide homozygosity except around the mating-type locus, our experiments provided one of the rare direct experimental evidence of sheltered load around mating-compatibility loci, which is crucial for our understanding of sex-related chromosome evolution.

### Competing interests

The authors have no competing interests.

## Acknowledgements

This work was supported by the EvolSexChrom ERC advanced grant #832352 (H2020 European Research Council) to T.G. We acknowledge the ImagoSeine core facility of the Institut Jacques Monod, member of the France BioImaging infrastructure (ANR-10-INBS-04) and GIS-IBiSA and the support of the Region Ile-de-France (Sesame). The funders had no role in study design, data collection and analysis, decision to publish, or preparation of the manuscript. We thank Philippe Silar for isolating heterokaryotic strains used here of *Podospora anserina*, *Schizothecium tetrasporum* and *S. tritetrasporum*, for designing the primer pairs, for the picture of spores in Figure 1 and for advice during the experiment design. We thank Asen Daskalov, Kamalraj Subban and Alexandra Granger-Farbos for isolating *MAT1-1* and *MAT1-2* homokaryons of the *Podospora anserina* strains used in this study. We thank Albane Thibeault for help with nucleus counts. We thank Jacqui Shykoff for discussions on statistical analyses. We thank the collectors of soil and animal faeces: Philippe Silar, Delphine Paumier, Yves Hurand, Pierre Defos du Rau, Jean-Marie Ouary (Milles Traces association), Jérome Letty (Office français de la biodiversité), Paul Jay, Stella McQueen, Karen Vanderwolf, Jacqui Shykoff, Lucas Bonometti and Pierre Gladieux.

## Author contribution

Conceptualization: TG, FEH and LG. Data curation: LG, SB, EDF. Formal analysis: LG, CL, EDF. Funding acquisition: TG. Investigation: LG, TG, FEH, SB, EC and EDF. Supervision: TG and FEH. Visualization: LG and EDF. Writing: LG, TG. Writing – review & editing: LG, TG, FEH, CL, EDF.

## Supplementary Figure legends

**Supplementary Figure 1:**
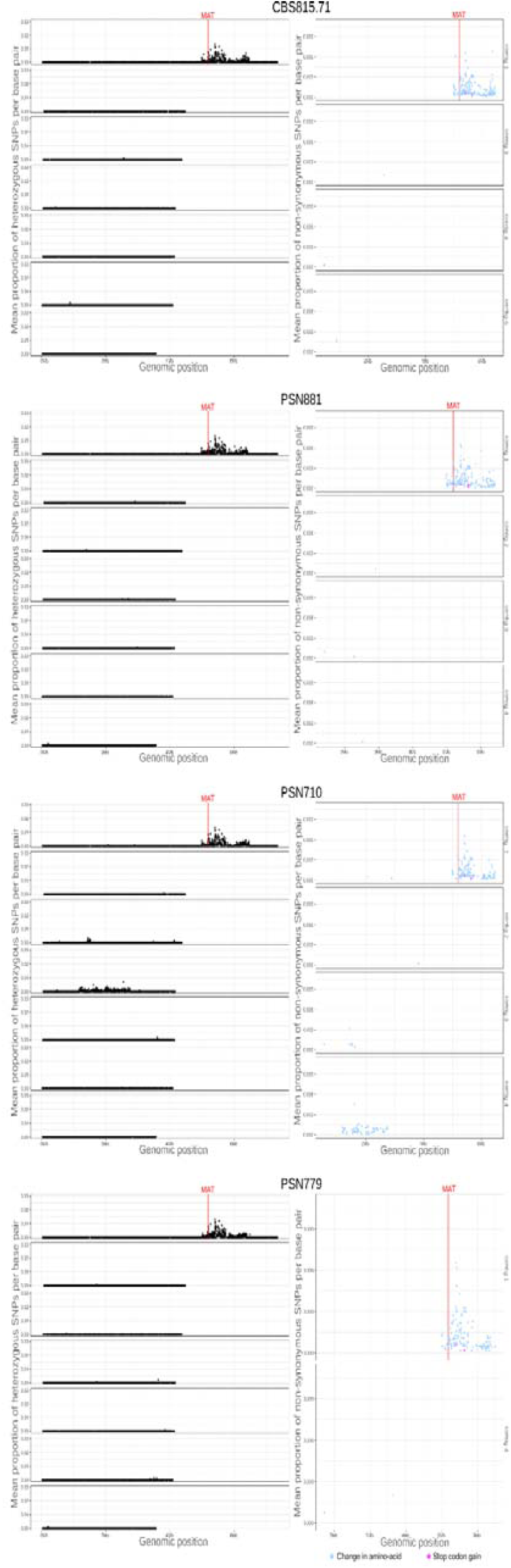
Genomic analyses of heterozygosity and mating-type specific putative deleterious mutations (non-synonymous SNPs) along the chromosomes of four strains of *Schizothecium tetrasporum*. The left column depicts the genome-wide per-gene density in heterozygous SNPs for the four *S. tetrasporum* strains for which genomes were available. Each dot represents one gene, the Y axis corresponding to the mean proportion of heterozygous SNPs per base pair for each gene and the X axis to the genomic position along the CBS815.71-sp3 assembly in megabases (Mb). The seven longest contigs in the assembly are represented, by length order. The right column depicts the genome-wide per-gene density in heterozygous non-synonymous SNPs. Each dot represents one gene, the Y axis corresponding to the mean proportion of heterozygous non-synonymous SNPs per base pair; these SNPs introduced either a missense mutation (i.e., a change in amino-acid; blue dot) or the gain of a stop codon (pink dot) compared to the reference genome CBS815.71-sp3. The X axis corresponds to the genomic position along the CBS815.71-sp3 assembly in megabases (Mb). Dots indicating a null value have been removed for clarity. The mating-type locus (MAT) position is indicated by a vertical red line.

**Supplementary Figure 2:**
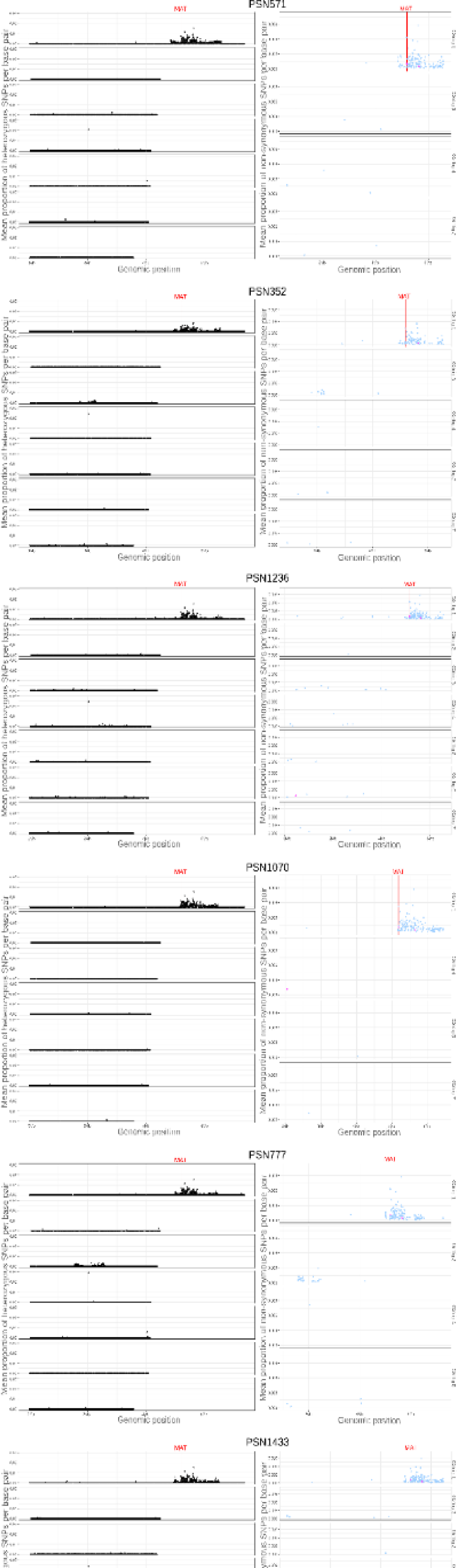
Genomic analyses of heterozygosity and mating-type specific possible deleterious mutations (non-synonymous SNPs) along the chromosomes of nine strains of *Schizothecium tritetrasporum*. The left column depicts the genome-wide per-gene density in heterozygous SNPs for the nine *S. tritetrasporum* strains for which genomes were available. Each dot represents one gene, the Y axis corresponding to the mean proportion of heterozygous SNPs per base pair for each gene and the X axis to the genomic position along the CBS815.71-sp3 assembly in megabases (Mb). The seven longest contigs in the assembly are represented, by length order. The right column depicts the genome-wide per-gene density in heterozygous non-synonymous SNPs. Each dot represents one gene, the Y axis corresponding to the mean proportion of heterozygous non-synonymous SNPs per base pair; these SNPs introduced either a missense mutation (i.e., a change in amino-acid; blue dot), the gain of a stop codon (pink dot) or the loss of a start codon (orange dot) compared to the reference genome CBS815.71-sp3. The X axis corresponds to the genomic position along the CBS815.71-sp3 assembly in megabases (Mb). Dots indicating a null value have been removed for clarity. The mating-type locus (MAT) position is indicated by a vertical red line.

**Supplementary Figure S3:**
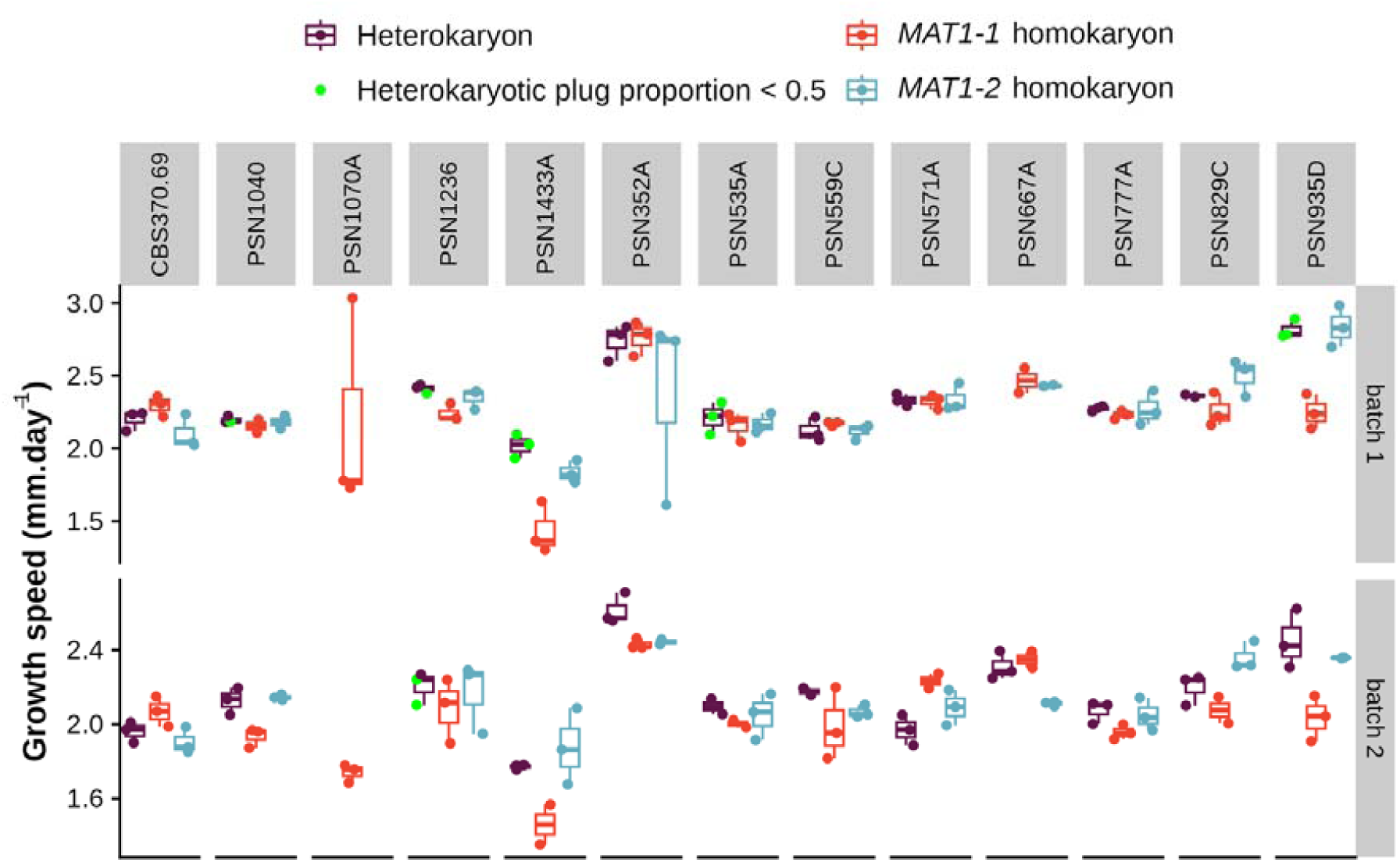
Mycelium growth speed of heterokaryons versus homokaryons for 13 strains of *Schizothecium tritetrasporum*. Mycelium growth speed (mm.dayL¹) of heterokaryons (in purple), *MAT1-1* homokaryons (in red) and *MAT1-2* homokaryons (in blue), of the three replicates for each of two batches, for the 13 *S. tritetrasporum* strains, between day 6 and day 14, grown under light conditions at 22°C. Green dots highlight the heterokaryon replicates for which fewer than 50% of the nine mycelium plugs from their Petri dish were genotyped heterokaryotic. Boxplot elements: central line: median, box limits: 25th and 75th percentiles, whiskers: 1.5× interquartile range.

**Supplementary Figure S4:**
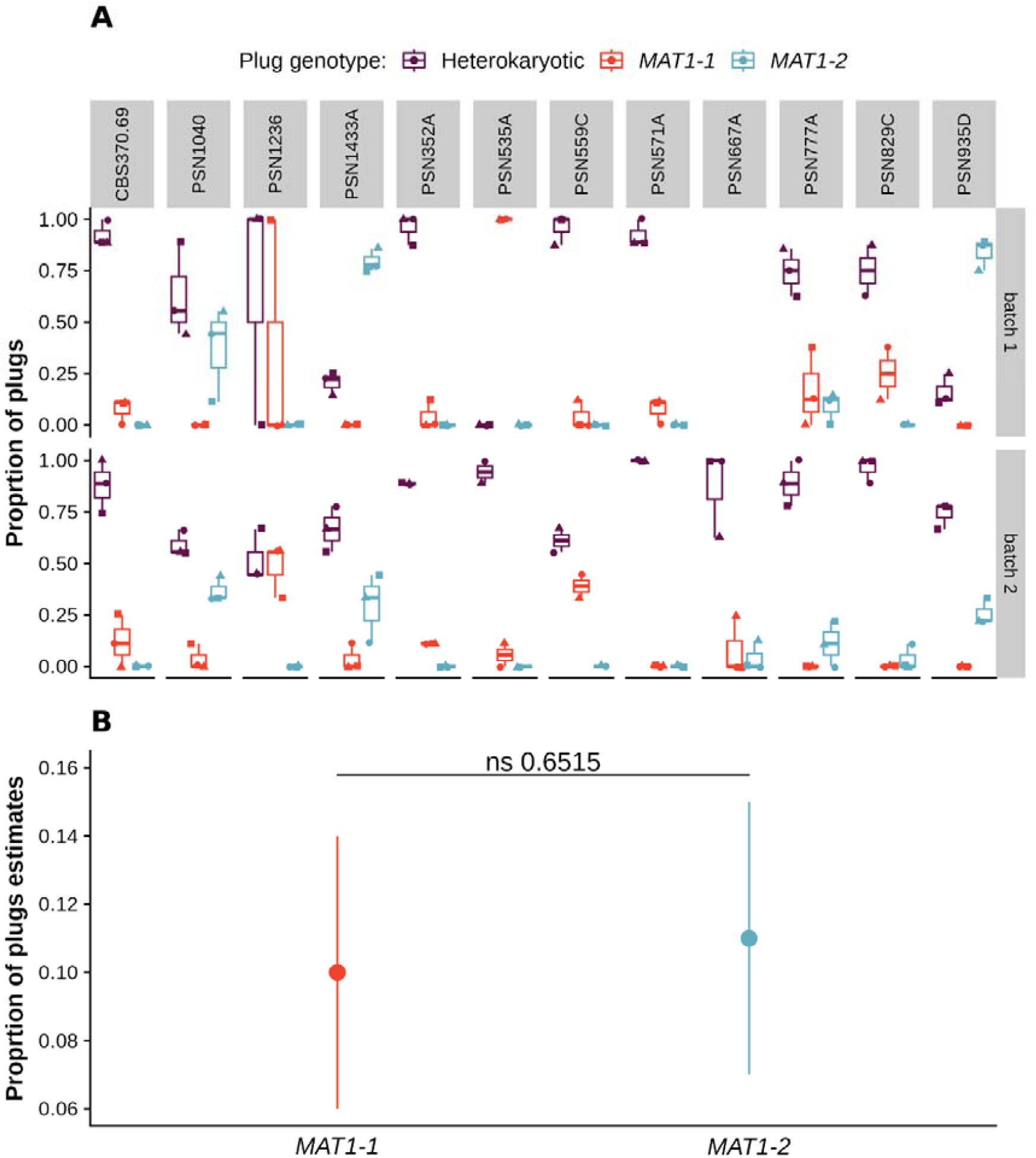
Nuclear loss in *Schizothecium tritetrasporum* heterokaryotic mycelium. A) Proportions of plugs with only *MAT1-1* (in red), only *MAT1-2* (in blue) or both MAT (in purple) detected in *S. tritetrasporum* heterokaryotic mycelium after 14 days of growth. Proportions were calculated out of nine plugs per plate that were genotyped by PCR in three replicates per strain for each of two batches. Symbol shapes correspond to replicate plates. Boxplot elements: central line: median, box limits: 25th and 75th percentiles, whiskers: 1.5× interquartile range. B) Model estimates of proportions of plug with only the *MAT1-1* or the *MAT1-2* allele detected by PCR in *S. tritetrasporum* heterokaryotic mycelium. Dots and bars represent marginal means and standard error estimated from a generalized linear mixed-effect model with a Binomial distribution modelling plug proportion with nuclear loss using the genotype of the plug after the loss (i.e. *MAT1-1* or *MAT1-2*) as a fixed effect and strain ID and batch as random effects. The p-value corresponds to a pairwise t-test (ns: non-significant).

**Supplementary Figure S5:**
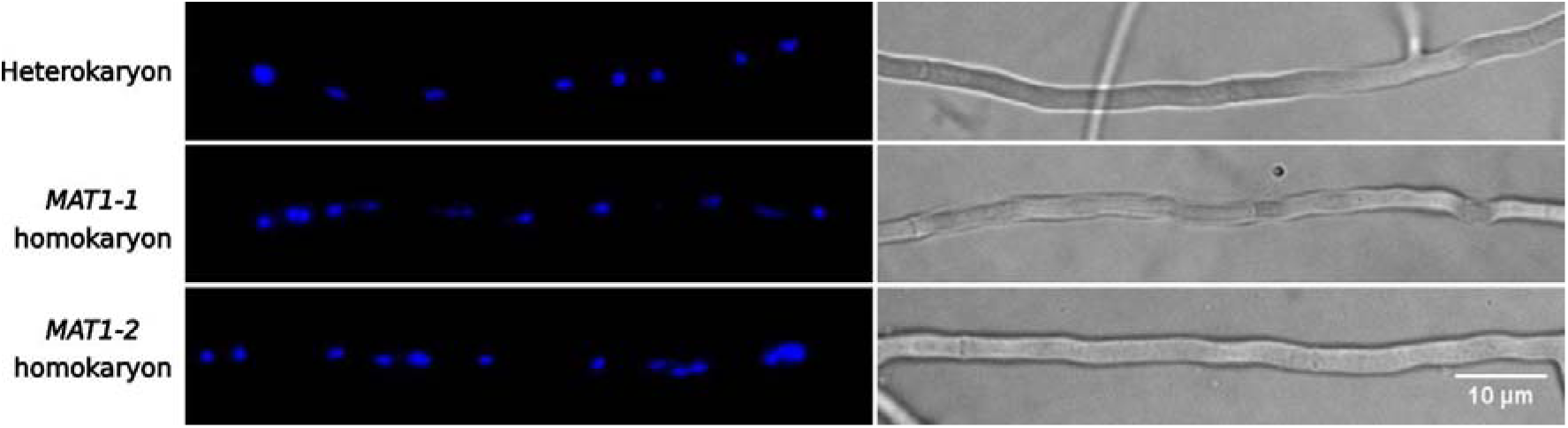
Distribution of nuclei in vegetative hyphae of the *Podospora anserina* S strain. Epifluorescence microscopy pictures of vegetative hyphae of the heterokaryon, the *MAT1-1* homokaryon and the *MAT1-2* homokaryon of the *P. anserina* S strain. Left column: DAPI stained nuclei, images correspond to z-projections of three z-slices separated by 1 µm. Right column: bright-field images corresponding to the slice in the middle of the z-stack. Arrow heads: septa. Scale bar: 10 µm.

**Supplementary Figure S6:**
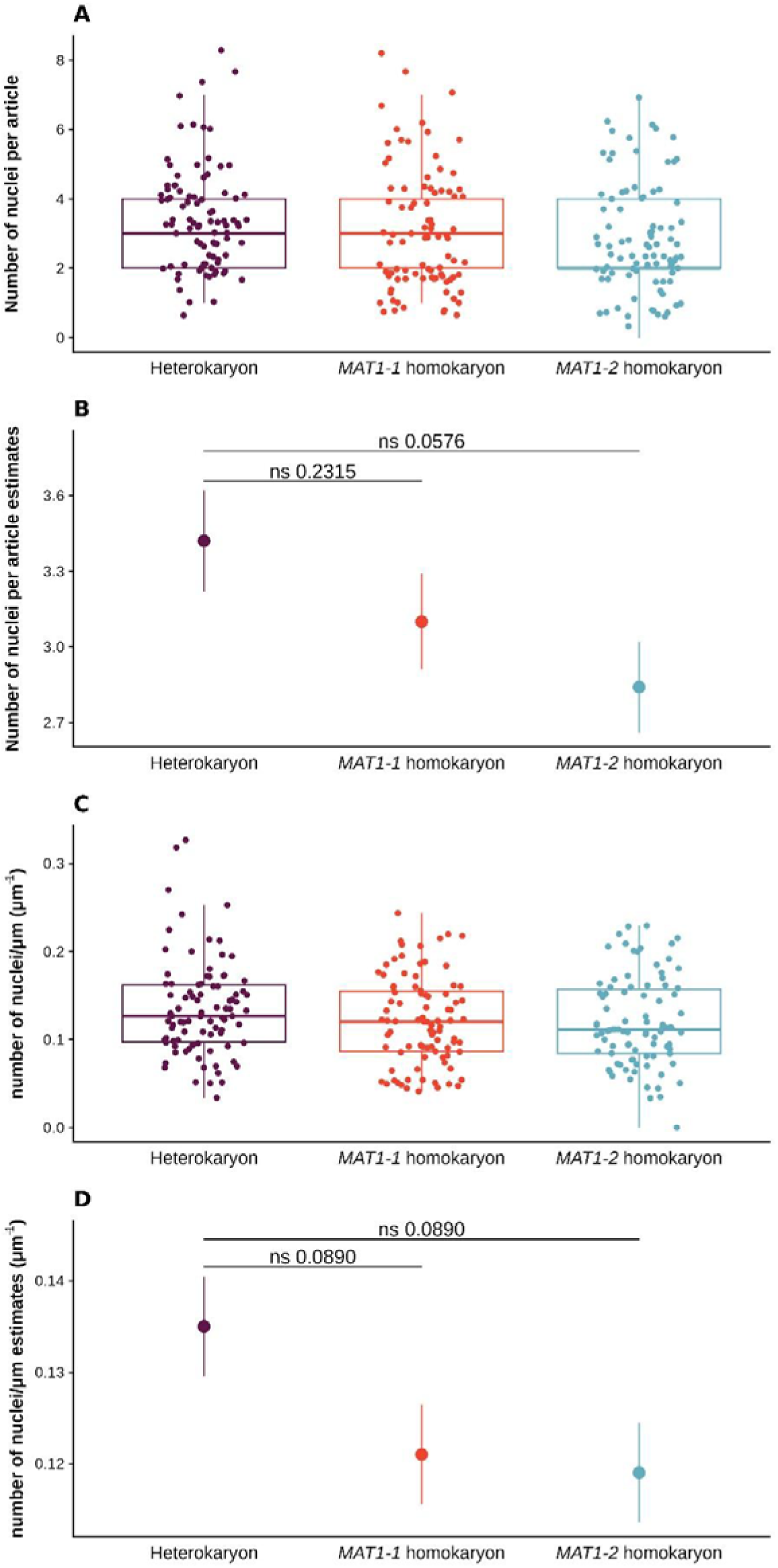
Number of nuclei and their density per article in heterokaryotic versus homokaryotic mycelia in the *Podospora anserina* S strain. A) Number of nuclei per article in heterokaryons (in purple), *MAT1-1* homokaryons (in red) and *MAT1-2* homokaryons (in blue), in 30 articles of three replicate Petri dishes of the *P. anserina* S strain at day 6 of growth, at 18°C under light. Boxplot elements: central line: median, box limits: 25th and 75th percentiles, whiskers: 1.5× interquartile range. B) Model estimates of nucleus number per article in heterokaryons (in purple), *MAT1-1* homokaryons (in red) and *MAT1-2* homokaryons (in blue) of the *P. anserina* S strain. Dots and bars represent marginal means and standard errors estimated from a generalised linear model with a Poisson distribution explaining nucleus number per article using nuclear status (i.e. *MAT1-1* homokaryon, *MAT1-2* homokaryon or heterokaryon) as fixed effects. P-values correspond to pairwise z-tests adjusted for multiple testing using the Holm method (ns: non-significant). **C)** Density of nuclei per article (nb of nuclei.μL¹) in heterokaryons (in purple), *MAT1-1* homokaryons (in red) and *MAT1-2* homokaryons (in blue), in 30 articles of three replicate Petri dishes of the *P. anserina* S strain at day 6 of growth, at 18°C under light. Boxplot elements: central line: median, box limits: 25th and 75th percentiles, whiskers: 1.5× interquartile range. **D)** Model estimates of the density of nuclei per article in heterokaryons (in purple), *MAT1-1* homokaryons (in red) and *MAT1-2* homokaryons (in blue) of the *P. anserina* S strain. Dots and bars represent marginal means and standard errors estimated from a linear model explaining nucleus density per article using nuclear status (i.e. *MAT1-1* homokaryon, *MAT1-2* homokaryon or heterokaryon) as fixed effects. P-values correspond to pairwise F-tests adjusted for multiple testing using the Holm method (ns: non-significant).

## Notes

### Competing Interest Statement

The authors have declared no competing interest.

### Summary of Updates

Addition of genomic analyses and text revision

